# The Ry_sto_ immune receptor recognizes a broadly conserved feature of potyviral coat proteins

**DOI:** 10.1101/2021.05.20.444601

**Authors:** Marta Grech-Baran, Kamil Witek, Jarosław Poznański, Anna Grupa-Urbańska, Tadeusz Malinowski, Małgorzata Lichocka, Jonathan DG Jones, Jacek Hennig

## Abstract

Potyviruses are the largest group of plant RNA viruses, causing significant losses in many crops. Among them, potato virus Y (PVY) is particularly important, and enhances the severity of infections by other viruses. The *Ry_sto_* gene confers PVY resistance and encodes a TIR-NLR intracellular immune receptors that recognizes PVY coat protein (CP). To define a minimal CP fragment sensed by Ry_sto_, we created a series of truncated CP variants and expressed these CP derivatives in *Ry_sto_* transgenic plants. Deletions that affect the 149 amino acid CP core region lose the ability to trigger *Ry_sto_*-dependent defence activation. Furthermore, point mutations in the amino acid residues Ser_126_, Arg_157_, and Asp_201_ of the highly conserved RNA-binding pocket of potyviral CP, reduce or abolish Ry_sto_-dependent responses, demonstrating that appropriate folding of the CP core is required for Ry_sto_-mediated recognition. Consistent with these data, we found Ry_sto_ recognises CPs of various viruses that share a similar core region, but not those lacking it. Finally, we demonstrated that Ry_sto_ provides immunity to plum pox virus and turnip mosaic virus, demonstrating its wide range of applications in disease-resistant crop engineering. In parallel, we showed that CP triggered Ry_sto_ activation is SAG101- but not PAD4- or SA-level dependent. Our findings shed new light on how R proteins can detect viruses by sensing highly conserved structural patterns.

## Introduction

The plant immune system is a multilayered detection and signalling network. To successfully restrict and thwart viruses, plants use RNA silencing, translation repression, and *Resistance* (*R*) gene-mediated mechanisms (Wersch et al., 2020; Leisner and Schoelz, 2018). Most plant *R* genes encode intracellular nucleotide-binding leucine-rich repeat (NLR) receptors. Some of these receptors carry an N-terminal TIR domain and are termed TNLs, whereas CNLs carry an N-terminal coiled-coil domain (Jones et al., 2016). A growing body of evidence suggests that in plants, as in mammals, NLR activation results from induced oligomerization of NLRs to form signalling-active scaffolds (Saur et al., 2020). Recent structural elucidation of the CNL: HOPZ-ACTIVATED RESISTANCE1 (ZAR1) and TNL-type: ROQ1 (recognition of XopQ 1) and RPP1 (Resistance to *Peronospora parasitica* 1) immune receptors provided new insights into plant immune receptor activation (Wang et al., 2019; Martin et el., 2020). ZAR1 is as an ADP-bound monomer in the resting state and after activation forms a pentameric “resistosome” (Wang et al., 2019). ROQ1 and RPP1, when activated, form tetrameric complex (Martin et al., 2020; Ma et al., 2020). Historically, pathogen gene products that are detected by matching NLR receptors were named avirulence (Avr) factors (Flor, 1956; Wersch et al., 2020). The activation of NLRs leads to a suite of downstream defence responses, frequently culminating in a hypersensitive cell death response (HR) at the infection site (Lolle et al., 2020). This model of resistance is often referred to as effector-triggered immunity (ETI) and involves NLR-dependent direct or indirect pathogen recognition (Zipfel, 2014). The majority of viral effectors is functionally represented by movement proteins, replicases, and coat proteins (CPs), with all playing various roles over the course of the infection (Leisner et al., 2018). Due to the small size of the viral genome, some of the proteins might be multifunctional, like the P6 protein of the cauliflower mosaic virus, which facilitates several steps of the infection, including modulation of host defences (Leisner and Schoelz, 2018). Despite progress made over the last decade, the relationship between the function of recognized viral proteins and their perception by NLRs still needs to be addressed.

Potyviruses are the largest genus of plant viruses infecting a broad range of dicot and monocot crops (Hoffius et al., 2007). Potato virus Y (PVY), plum pox virus (PPV), turnip mosaic virus (TuMV), papaya ringspot virus; (PRSV), soybean mosaic virus (SMV), sweet potato feathery mottle virus (SPFMV) and others cause billions of dollars worth of annual losses worldwide (da Silva et al., 2020). One of the most extensively studied, potato virus Y (PVY) is the most damaging viral pathogen of Solanaceae crops (da Silva et al., 2020). Infective PVY virions are flexuous filaments where coat protein (CP) subunits assemble in a helical mode and are bound to a monopartite positive-sense single-stranded (ssRNA) genome (Zamora et al., 2017). The CP, apart from its principal role as a structural protein of the virion, is also involved in transmission by aphids, and movement and regulation of viral RNA amplification (Revers et al., 2015).

Several mechanisms for the perception of viruses by R proteins have been proposed. Some NLR proteins may recognise more than one effector from various pathogens. For instance, Rx, a resistance protein against potato virus X (PVX), sensed the conserved CP motifs from various potexviruses, except the PVX_HB_ strain (Baures et al., 2008). Similarly, the N protein that confers resistance against the tobacco mosaic virus (TMV) via recognition of the helicase domain of TMV replicase (p50), can perceive several tobamoviruses (Whitham et al., 1994). Next, *R* gene loci carrying allelic variants encoding proteins may also recognize distinct effectors (Le Roux et al., 2015). Likewise, L proteins, which recognise tobamoviral CPs (Cooley et al., 2000; Tomita et al., 2008), and HRT or RCY-1, at the Arabidopsis RPP8 locus, confer resistance to turnip crinkle virus (TCV) and cucumber mosaic virus (CMV), respectively (Takahashi et al., 2002). In turn, the *Magnaporthe oryzae* effectors Avr-Pia, Avr1-CO39, Avr-PikD, and AvrPiz-t, despite being sequence-unrelated, share the same architecture and all bind to the rice resistance protein Pik-1 via HMA domains (Bialas et al., 2020; Maqbool et al., 2015). Similarly, five AvrPm3 effector proteins from wheat powdery mildew form a large group of proteins with low sequence homology but structural similarity that are perceived by the wheat NLR receptor PM3 (Bourras et al., 2019). Nevertheless, the question of whether the way the receptor recognizes Avr influences the range of its mediated immunity remains to be fully addressed.

We previously showed that the TNL immune receptor Ry_sto_ perceives the PVY CP and is required for PVY-triggered immunity (Grech-Baran et al., 2020). *Ry_sto_* confers an extreme resistance response in potato that prevents viral replication without triggering cell death. Nonetheless, overexpression of the PVY CP in *Ry_sto_*-expressing plants can trigger a macroscopic HR in heterologous systems, such as *Nicotiana tabacum*. To date, the molecular mechanism of *Ry_sto_*-triggered immunity is largely unknown. We show here that Ry_sto_ perceives a specific molecular pattern that is similar in multiple potyviruses. Hence, it may serve as a rapid selectable marker in Ry_sto_-conferred immunity. Furthermore, we revealed that Ry_sto_ might be used to develop durable potyviral disease resistance. In parallel, we shed light on new elements of the signalling mode of Ry_sto_-mediated immune response.

## Results

### The core region of CP is both necessary and sufficient for eliciting cell death in *Ry_sto_* transgenic plants

To discover the molecular mechanisms underlying PVY CP perception by the Ry_sto_ protein, a PVY CP structural model (Protein Data Bank 6HXX) was analysed using the YASARA structure package (Krieger and Vriend, 2014). A single PVY CP unit consists of 267 amino acids with a globular core subdomain defined from Q_77_ to K_226_ that forms seven α-helix folds and a β-hairpin. This central domain is flanked by putative disordered regions comprising 76 and 41 amino acid residues at the N- and C-termini, respectively (Fig. 1a, Suppl. Fig. 1). To determine which CP region is recognised by Ry_sto_, the truncated CP variants CP_77-267_, CP_1-226_, and CP_77-226_, as well as the full CP, were cloned and expressed in *Ry_sto_ Nicotiana tabacum* and control, non-transgenic plants (Fig. 1a). Neither the N-nor C-terminus of PVY CP was required to elicit the response in *Ry_sto_ N. tabacum*, because the CP_77-226_ mutant still induced cell death (Fig. 1b). Based on the *in silico* model, we removed two amino acids from each end of the CP (CP_79-226_ or CP_77-224_) because this was predicted to disrupt the distal α-helices and likely hamper recognition by distorting the CP structure. Neither of these truncated constructs could elicit cell death in *Ry_sto_ N. tabacum* (Fig. 1b, c), indicating that the full-length CP core region is required for perception by Ry_sto_.

**Fig. 1.**
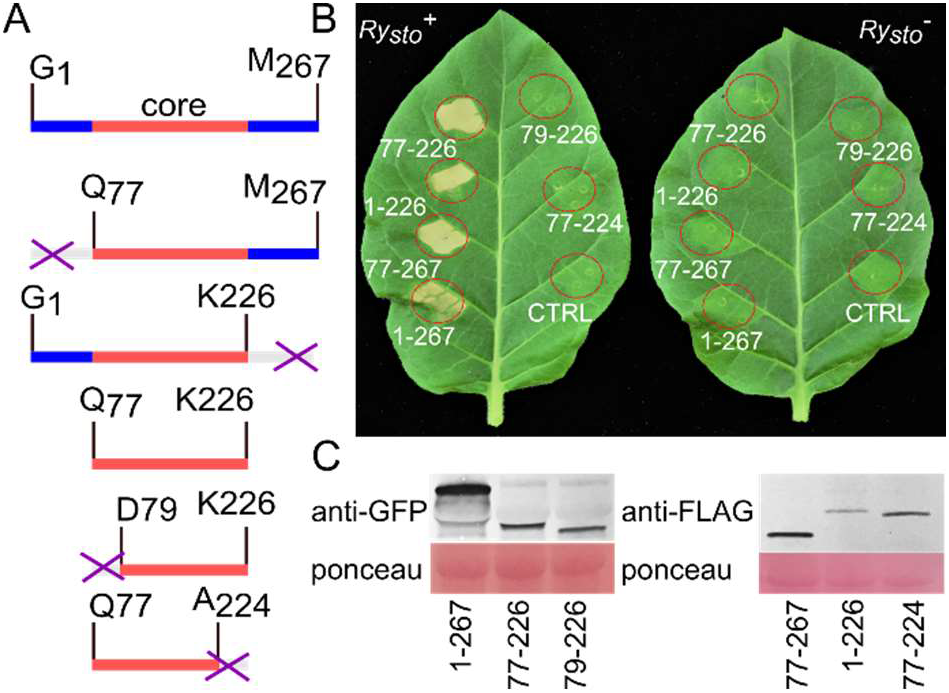
Mapping of the minimal CP fragment perceived by Ry_s_to. (a) Schematic representation of PVY CP and corresponding truncations. Blue, unstructured N- and C-terminal regions; pink, the core subdomain; white and crossed out, the deleted fragments. (b) Developments of cell death in leaves of *Ry_sto_* or control tobacco plants infiltrated with *Agrobacterium* expressing full-length CP or its truncated variants. As a negative control (CTRL) empty vector was used. Photographs were taken at 3 days postinfiltration (dpi). (c) *In planta* accumulation of CP variants. For anti-FLAG, and anti-GFP immunoblots of the full-length CP or its truncated variants, total proteins were prepared from the leaves tested at 3 days after agroinfiltration. Staining of RuBisCo with Ponceau S was used as a loading control.

### Conserved putative core CP residues at the protein-RNA interface are required for eliciting cell death in *Ry_sto_ N. tabacum* plants

The cryo-EM model of PVY (Kežar et al., 2019) suggested that the structure of PVY virions is stabilised by protein-ssRNA interactions involving the conserved residues S_125_, R_157_, and D_201_ at the ssRNA binding site of the CP core (Fig. 2a). Because the universally conserved RNA-binding pocket might be essential for capsid assembly and viral genome packaging (Zamora et al., 2017), we decided to investigate the role of the conserved residues in Ry_sto_-mediated CP core detection. Single (S_125_A, R_157_D, and D_201_R), double (S_125_A_R_157_D and S_125_A_D_201_R), or triple (S_125_A_R_157_D_D_201_R) amino acid substitutions were generated using site-directed mutagenesis. Following the transient expression of PVY CP mutants in *Ry_sto_* plants, three phenotypes were observed. A typical cell death phenotype was observed for the single substitutions S_125_A and R_157_D, whereas the R_157_D mutant and each of the double variants partially retained CP recognition, and triggered weaker cell death. In contrast, no cell death was observed when the triple mutant was tested (Fig. 2b). Expression of all proteins was confirmed by western blot analysis (Fig. 2c).

**Fig. 2.**
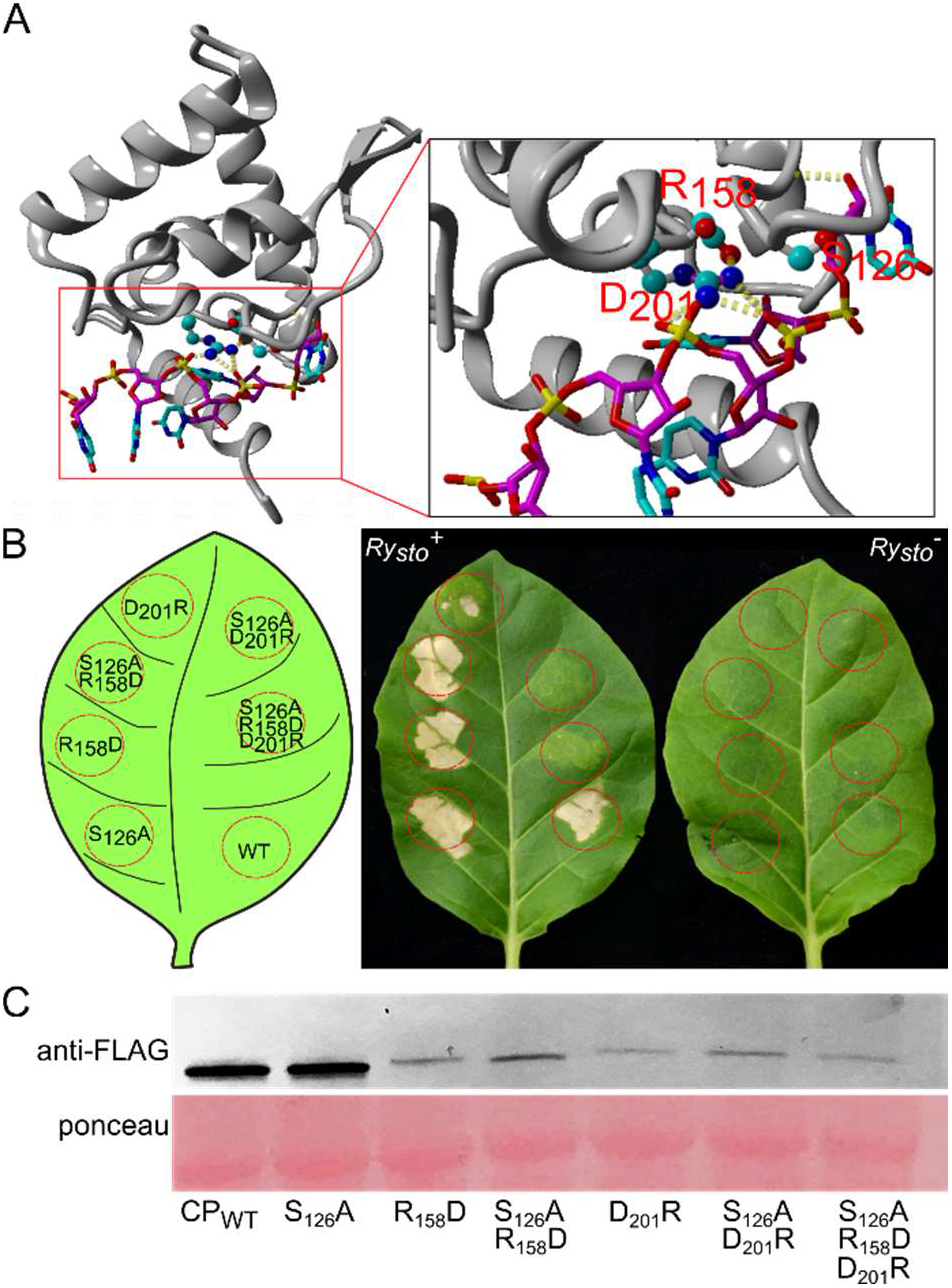
Effect of the conserved residues of protein-RNA interaction pocket site on Ry_sto_-mediated CP perception. (a) PVY CP core structural model. Inset: a detailed view of the highly conserved residues S_125_, R_157_, and D_201_ involved in PVY ssRNA binding. Electrostatic interactions are shown as yellow dashed lines. (b) Cell death phenotypes in leaves of *Ry_sto_* transgenic or control tobacco plants infiltrated with *Agrobacterium* expressing CP or mutated CPs with a single, double, or triple amino acid substitution. Photographs were taken at 3 days post-infiltration (dpi). (c) *In planta* accumulation of the CP mutant proteins. For anti-FLAG immunoblots of the full-length CP (positive control) and CP variants, total proteins were prepared from the leaves tested at 3 days after agroinfiltration. Staining of RuBisCo with Ponceau S was used as a loading control.

To quantify the Ry_sto_-mediated cell death response to the CP mutants, all CP versions were transiently expressed in Ry_sto_-expressing tobacco leaves and electrolyte leakage was measured at 18, 21, 24, 27, and 30 h after infiltration (hpi). Full-length PVY CP was used as a control. All single and double mutants followed the conductivity pattern of the wild-type (WT) CP, with the maximum conductivity detected at 27 hpi, although the ion leakage was lower than that of the control at every time point, especially in the single and double mutants with the D_201_R mutation. As expected, the triple S_125_A_R_157_D_D_201_R mutant activated no ion leakage (Fig. 3a). This supports the hypothesis that the S_125_, R_157_, and D_201_ pocket is essential for proper folding of the PVY CP central core region and, consequently, for Ry_sto_-mediated recognition.

**Fig. 3.**
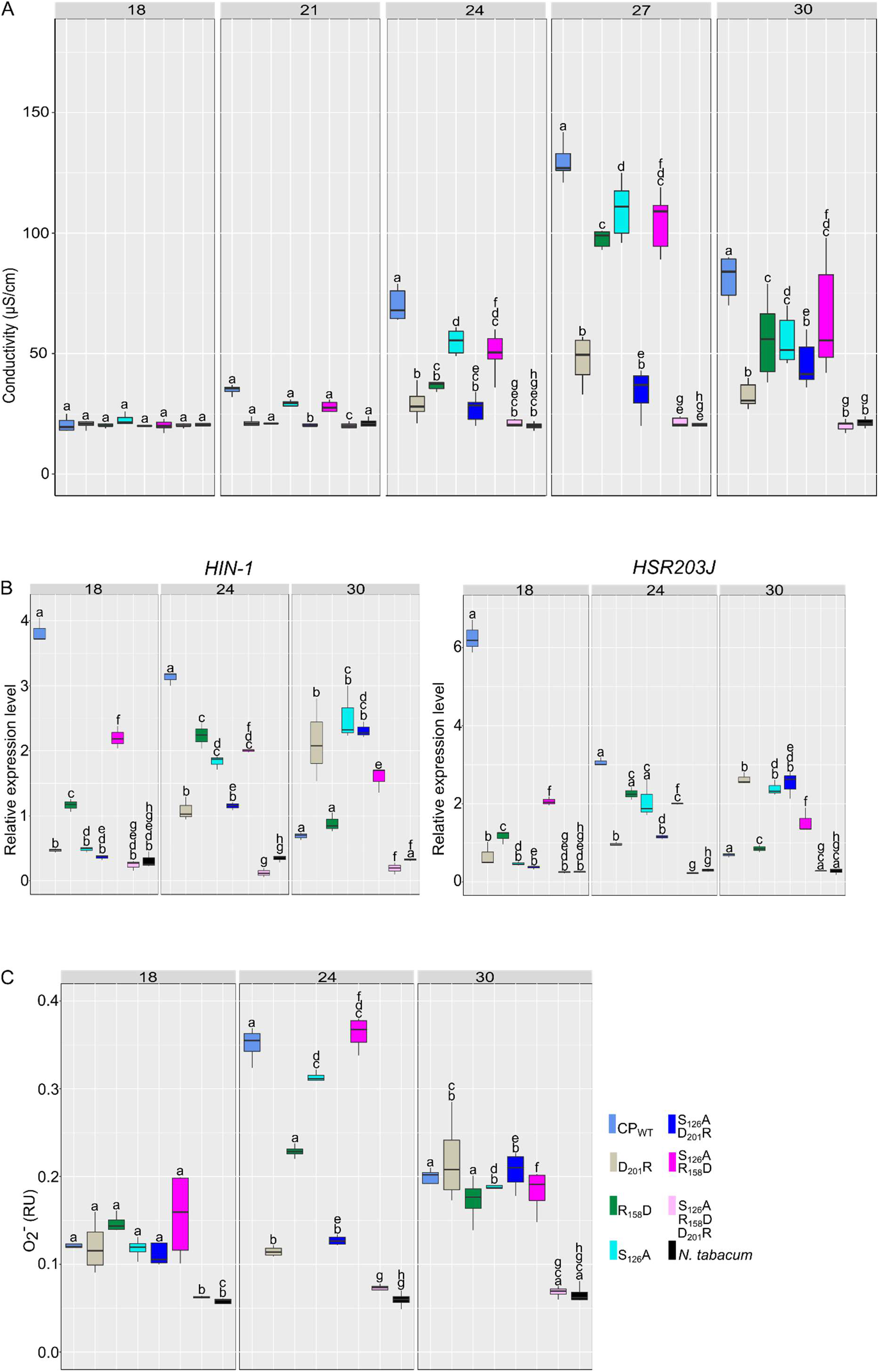
Mutations of the conserved residues of CP core are associated with a delayed or abolished onset of *Ry_sto_* detection. (a) Cell death measured by ion leakage in leaf discs from *Ry_sto_* transgenic tobacco leaves infiltrated with *Agrobacterium* carrying CP or its indicated variants. As a negative control nontransgenic *N. tabacum* infiltrated with the full-length CP was used. The electroconductivity was measured at 3 h intervals from 18 to 30 hours post-infiltration (hpi); *n* = 8. (b) Quantitative RT-PCR analysis of the defence response marker genes. *HIN-1* and *HSR203J* transcript accumulation was measured at 18, 24, and 30 hpi. *EF1* and *L23* expression were used as references. (c) ROS burst measured by the concentration of superoxide anion (O_2_^-^) detected in tested leaf discs in the presence of 0.05% nitroblue tetrazolium (w/v). Samples were collected every 3 h. O_2_^-^ content is expressed in relative units (RU = A_560_/6.3 cm^2^); *n* = 8. Data from each experiment were analysed using a repeated-measures ANOVA followed by Tukey’s HSD post-hoc test performed separately for each time point. Letters correspond to statistically homogeneous groups (*p* < 0.001).

The expression of defence-related marker genes *HSR203J* and *HIN-1* (Chichkova et al., 2004) was tested by quantitative polymerase chain reaction (qPCR) at 18, 24, and 30 hpi. In Ry_sto_ tobacco infiltrated with WT CP, the steady-state levels of both genes were at a maximum at 18 hpi and gradually decreased. All single and double mutants showed a significant decrease and delay in *HSR203J* and *HIN-1* expression. The observed changes were less pronounced in R_157_D and S_125_A/R_157_D in comparison with other mutated CP versions (Fig. 3b). Consistent with the observed phenotype, the triple mutant was unable to induce *HSR203J* and *HIN-1* expression (Fig. 3b).

Finally, ROS production following Ry_sto_ elicitation was measured. Surprisingly, the level of O_2_^-^ at 18 hpi was higher in R_157_D and S_125_A_R_157_D than in WT CP (Fig. 3c). Similarly, at 24 hpi, it remained elevated in comparison to that in WT CP or other mutants, which exhibited an impaired oxidative burst. The triple mutant did not activate an oxidative burst at any of the tested time points (Fig. 3c).

Taken together, these results suggest that mutations in the highly conserved amino acids of the CP core impaired or abolished the cell death phenotype in *Ry_sto_* plants by disturbing the three-dimensional structure.

### The EDS1-SAG101-NRG1 node is indispensable for Ry_sto_ CP triggered immunity

TNLs utilise a network of helper NLRs for a fully effective immune response. Two distinct complexes (NRG1/EDS1/SAG101 and ADR1/EDS1/PAD4) are required to execute TNL-initiated immunity (Sun et al., 2020); Ry_sto_ is no exception (Grech-Baran et al., 2020). To further investigate whether PAD4 or SAG101 are required for Ry_sto_-CP triggered response we transiently co-expressed Ry_sto_ with PVY CP or its triple inactive mutant in *pad4* or *pad4/sag101b N. benthamiana* and then quantified the HR (Fig. 4, Suppl. Fig. 8; Gantner et al., 2019). Ry_sto_-CP co-expression in *pad4* and WT *N. benthamiana* resulted in clear HR, while double *pad4/sag101b* knockouts remained symptomless (Fig. 4a). The triple CP mutant did not trigger any cell-death symptoms regardless of the background (Fig. 4a, b). Although, we observed some small reduction in the size of HR lesions in the *pad4* background, as compared to WT (Fig. 4b), it did not alter local Ry_sto_-CP resistance response activation. Thus, we conclude that SAG101 but not PAD4 is necessary for CP-triggered Ry_sto_ HR.

**Fig. 4.**
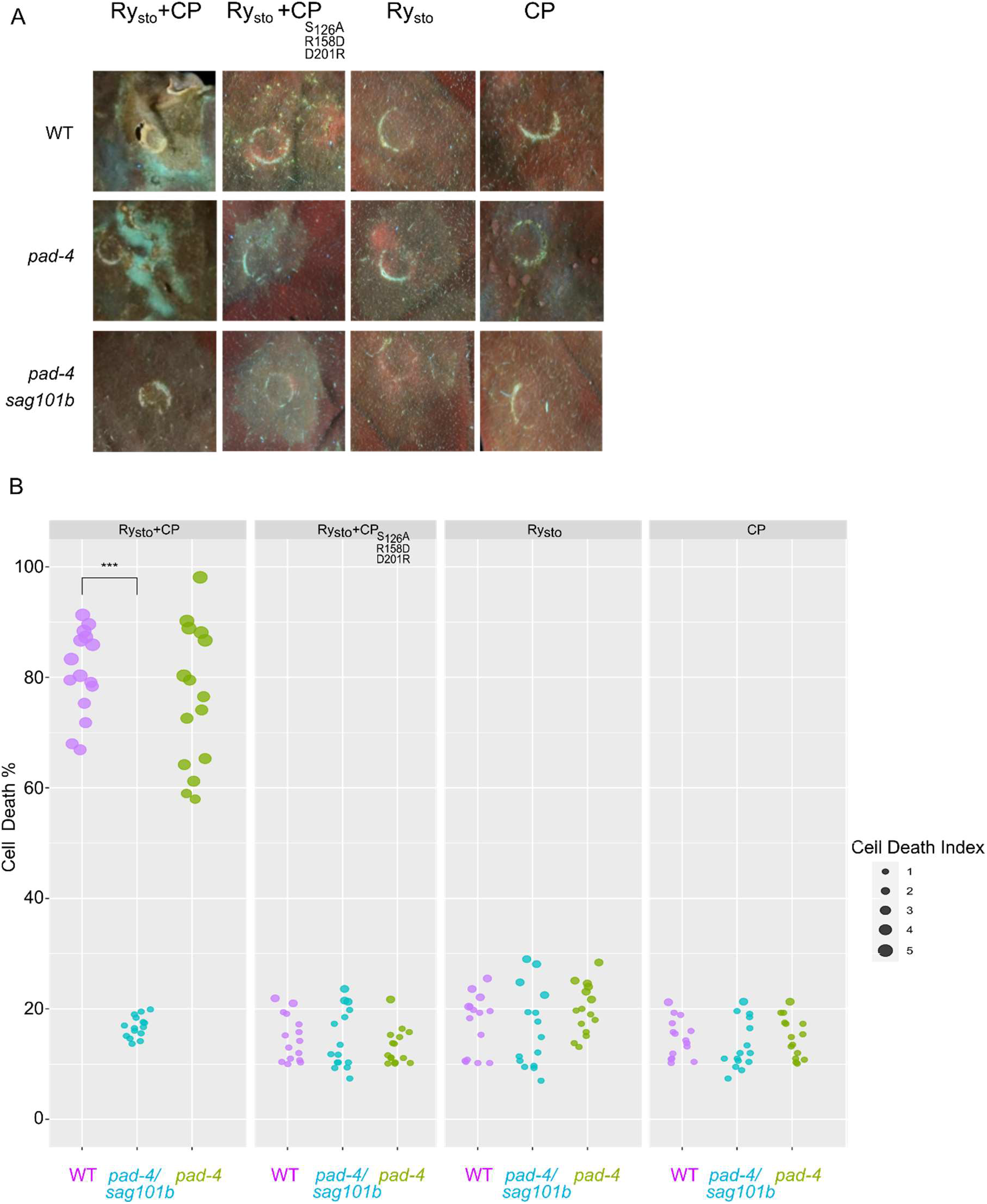
Ry_sto_-mediated immune response towards PVY CP is abolished in *sag101b* but not in *pad4* knockout lines. (a) Representative images of cell death phenotype after transient co-expression of the Ry_sto_ with PVY CP or Ry_sto_ with CP triple mutant in *pad-4*, or *pad-4*/*sag101b N. benthamiana* knockouts alongside with the wild-type control. The leaves were photographed five days after infiltration under UV light. CP triple mutant was used as a negative control. Cell death was scored five days after agroinfiltration (b). The results are presented as dot plots where the colour of a dot is proportional to the number of samples with the same score (count) within the same biological replicate. The statistical analysis was performed using ANOVA with post-hoc Dunnett’s test. The experiment was independently repeated at least three times.

### SA activation but not accumulation is required for CP-triggered immunity in *Ry_sto_* plants

SA is a crucial signalling molecule in the activation of plant immunity (Durner et al., 1997). To determine whether SA is involved in the initiation of the local Ry_sto_-mediated response, free SA levels were measured in *Ry_sto_* tobacco plants after overexpression of WT CP or the inactive mutant at different time points (18, 24 and 30 hpi) and temperatures (22 and 30°C). SA levels were significantly elevated from 24 hpi in leaves expressing CP at both temperatures, as compared to the mutant variant and mock-treated plants (Fig. 5a). Interestingly, the SA increase was less pronounced at 30°C (Fig. 5a). We have shown that transiently expressed *Ry_sto_* can trigger cell death in PVY-infected SA-deficient (*NahG*) *N. benthamiana* plants (Grech-Baran et al., 2020). To further explore the role of SA in Ry_sto_-triggered immunity, CP and Ry_sto_ were co-expressed in *NahG* or control tobacco plants. SA-deficient plants developed cell death symptoms similar to that of the controls (Fig. 5b). Furthermore, no significant differences in plasma membrane integrity or macroscopic tissue collapse were observed (Fig. 5c). These results suggest that even though the PVY CP triggers SA accumulation in Ry_sto_-expressing tobacco plants, the Ry_sto_-CP-related cell death is independent of SA accumulation. Similar results were obtained in *N. benthamiana* plants (Suppl. Fig. 2).

**Fig. 5.**
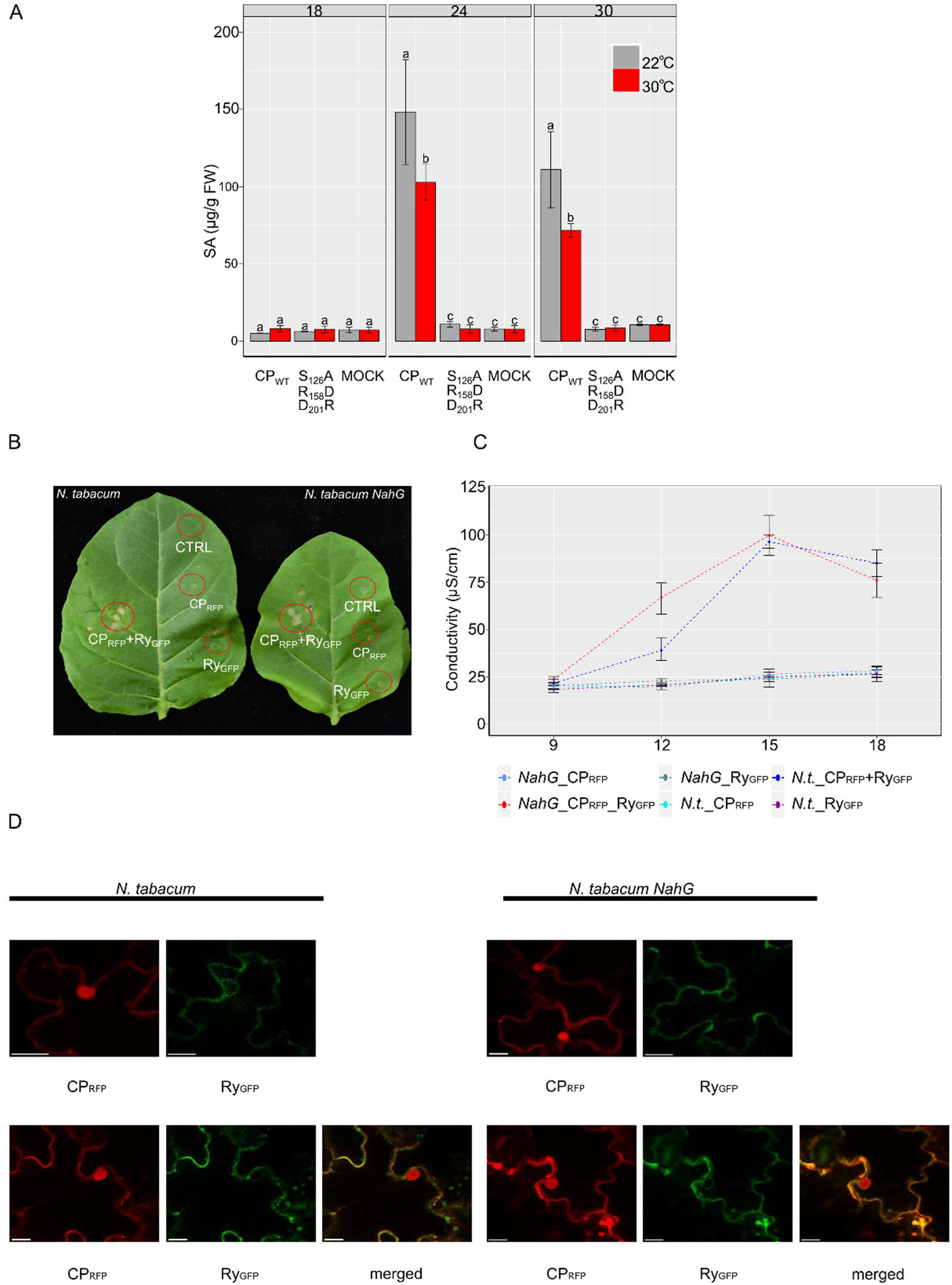
Role of SA in CP-triggered immunity in *Ry_sto_* plants. (a) Changes in free SA levels in *Ry_sto_* transgenic plants in response to CP under standard or elevated temperature conditions. Expression of the triple CP mutant or mock inoculation were used as a negative and positive control, respectively. At selected time points, samples of the plants tested were harvested and SA levels were measured by HPLC. Statistical analysis was performed using a one-way ANOVA with Tuckey’s HSD test (*p* < 0,01). (b) Ry_sto_ and CP co-expression results in tissue collapse in SA-deficient plants. Leaves of control or *NahG. tabacum* plants were infiltrated with *Agrobacterium* expressing Ry_sto_ or CP, or a combination of both. The photograph was obtained at 72 hours post-infiltration (hpi). Empty vector was used as a control. Expression of the *NahG* transcript was confirmed via semi-quantitative RT-PCR (Suppl. Fig. 7). (c) Ion leakage showing no differences in cell membrane permeability between *NahG* and control tobacco after CP and Ry_sto_ co-expression. Statistical analysis was performed using Student’s *t*-test. (d) Cellular localisation of CP and Ry_sto_ in representative *NahG* or control tobacco leaf epidermal cells transiently expressing CP, Ry_sto_, or both. Confocal images were obtained 72 hpi. For each variant, approximately 50 transformed cells were examined. Bars = 10 μm.

### Ry_sto_ recognises a range of CPs from multiple potyviruses

To determine whether CP of PVY shares structural similarities with other members of the *Potyvirus* genus, we constructed a phylogenetic tree of CPs from 65 potyviruses, which revealed a considerable divergence in their amino acid sequences (Fig. 6a, Suppl. Fig. 3). Despite the low similarity at their N- and C-termini, the central subdomain and the amino acids predicted to interact with ssRNA are conserved among all studied potyviruses (Fig. 6b, Suppl. Fig. 4, 5).

**Fig. 6.**
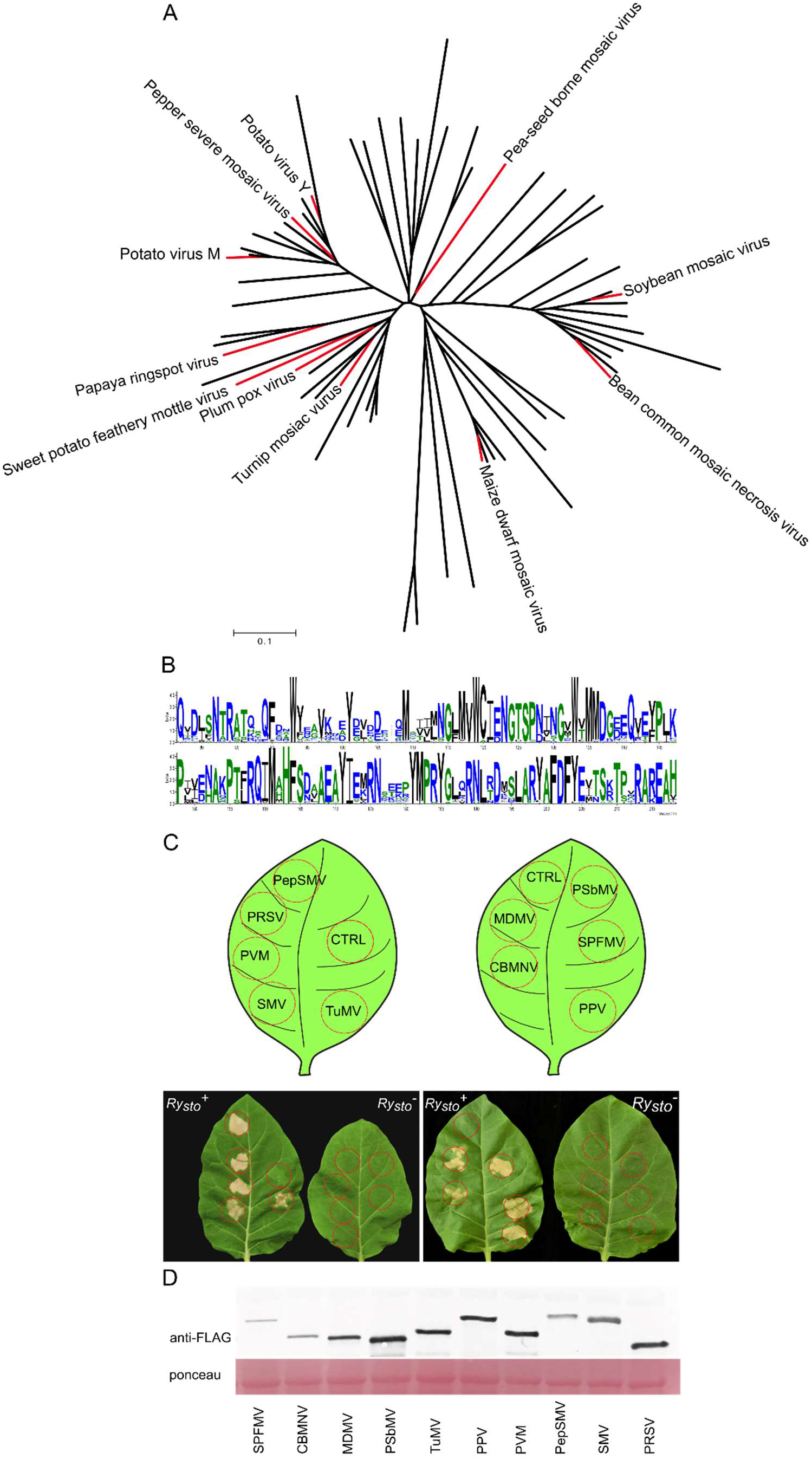
Coat proteins of multiple potyviruses that share a conserved structure trigger Ry_sto_-mediated immunity. (a) Phylogeny of *Potyviridae* CPs based on the amino acid sequences aligned using ClustalW and calculated with Mega7. Cloned sequences are marked in red. (b) Sequence logo analysis of CPs of 65 potyviruses depicting the conservation of the predicted core region, including the S_126__R_157__D_201_ ssRNA pocket binding site. (c) CPs of various potyviruses trigger Ry_sto_-mediated immunity. *Ry_sto_* or control tobacco plants were agroinfiltrated with multiple potyviral CPs: PRSV, papaya ringspot virus; PVM, potato virus M; SMV, soybean mosaic virus; PepSMV, pepper severe mosaic virus; CBMNV, bean common mosaic necrosis virus; MDMV, maize dwarf mosaic virus; TuMV, turnip mosaic virus; PSbMV, pea-seed born mosaic virus; SPFMV, sweet potato feathery mottle virus; or PPV, plum pox virus. All of them elicit Ry_sto_-dependent immune response. The photographs were obtained at 72 hours postinfiltration (hpi). Empty vector was used as a negative control (CTRL). (d) *In planta* accumulation of the potyviral CPs. For anti-FLAG immunoblots of viral CPs total protein extracts were prepared from leaves tested at 3 days after agroinfiltration. Staining of RuBisCo with Ponceau S was used as a loading control.

To test whether the structural similarities of CPs result in their recognition by Ry_sto_, we selected 10 economically important potyviruses from distantly related clades: PRSV, papaya ringspot virus; PVM, potato virus M; SMV, soybean mosaic virus; PepSMV, pepper severe mosaic virus; BCMNV, bean common mosaic necrosis virus; MDMV, maize dwarf mosaic virus; SCMV, turnip mosaic virus; TuMV, pea-seed born mosaic virus; SPFMV, sweet potato feathery mottle virus and PPV, plum pox virus. The sequences encoding these viral CPs were cloned and transiently expressed in transgenic Ry_sto_ or control tobacco plants. Within three days, cell death development was observed in Ry_sto_ plants for all tested CPs, whereas the controls remained symptomless (Fig. 6c). The expression of the viral proteins was confirmed by immunoblotting (Fig. 6d).

These results demonstrate that Ry_sto_ has a broad recognition spectrum and thus has considerable agronomic potential to control a number of economically significant potyviruses.

### A conserved potyviral CP core architecture is necessary for Ry_sto_-mediated recognition

To define the range of the Ry_sto_ mediated viral CP recognition, we tested the CP of ugandan cassava brown streak virus (CBSV; FJ185044.1) from *Ipomovirus* genus of the Potyviridae family (Monger et al., 2001). Structural modelling of CBSV CP revealed its low homology to the PVY CP structure, with significant differences in the central core region (Fig. 7a, b). Since CBSV CP does not have the strictly conserved LARY (A/G) FDFYE sequence, it cannot form the proper RNA binding pocket structure, resulting in a different core folding pattern (Fig. 2a, Fig. 7a). We tested whether this might result in lack of recognizability by Ry_sto_ (Fig. 7c) by cloning CBSV CP and transiently expressing it in transgenic Ry_sto_ and control tobacco plants. Although we could detect CBSV CP accumulation, no cell death was observed (Fig. 7c, d), consistent with the interpretation that recognition is dependent on the highly conserved CP core architecture in PVY and its closer relatives.

**Fig. 7.**
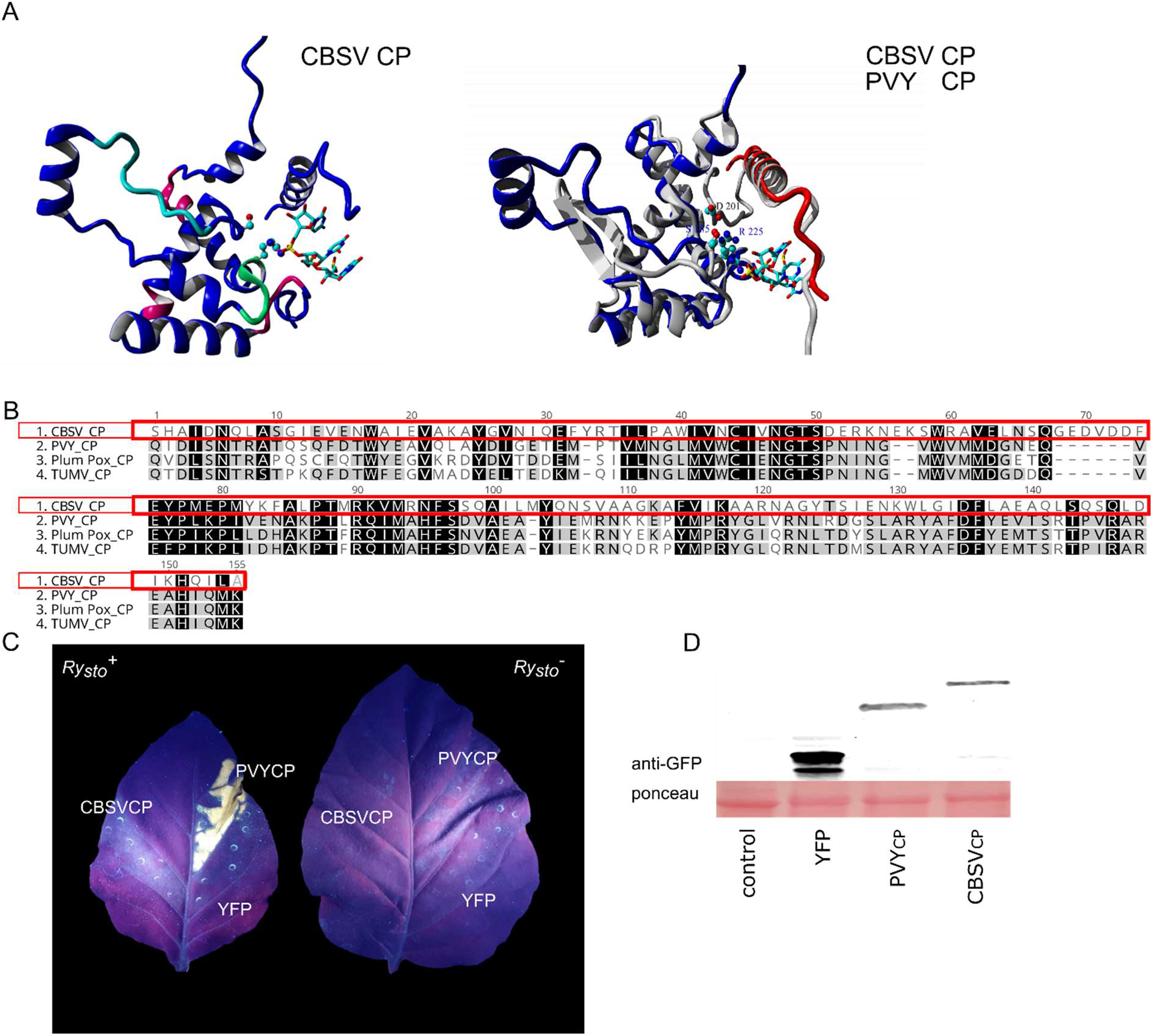
Variation in potyviral CPs core domain architecture results in lack of recognition by Ry_sto_. (a) The overall structure of CBSV CP (left) and structural alignment of CBSV and PVY CPs (right). The CBSV CP’s N-terminus - blue, C-terminus - red and PVY CP structure – grey (b) Amino acid sequence alignment of the predicted core regions of CPs from representative potyviruses recognised by Ry_sto_ *vs*. CBSV (red frame) (c) A representative image showing that CBSV CP is unable to elicit cell death in Ry_sto_-expressing plants. *Ry_sto_* transgenic and control tobacco plants were infiltrated with *Agrobacterium* expressing PVY CP-GFP, CBSV CP-GFP, or YFP alone as a control. Cell death development was observed only in *Ry_sto_* plants for PVY CP. The image was taken under UV light (365nm). (d) *In planta* accumulation of the viral CPs. Total protein extract was obtained from Ry_sto_-expressing plants at 72 hpi. Staining of RuBisCo with Ponceau S was used as a loading control.

### *Ry_sto_* restricts the systemic spread of PPV and TuMV

To test whether Ry_sto_ mediated recognition of the range of potyviral CPs also enables resistance against potyviruses other than PVY, we chose two economically important viruses, PPV and TuMV. PPV, a causative agent of Sharka disease, is the most important viral disease of stone fruit crops worldwide (Maejima et al., 2020), while TuMV damages various *Brassicacae* and other crops (Palukaitis and Kim, 2021).

Although PPV is unable to infect tobacco systemically, it can replicate in inoculated leaves (Saenz et al., 2002). We site-inoculated leaves of transgenic *Ry_sto_* and control tobacco plants with a severe PPV isolate (DSMZ No.: PV-0001). Seven days after infection, *Ry_sto_* plants exhibited a local cell death response, while no HR was detected in controls (Fig. 8a). To investigate the specificity of Ry_*sto*_-mediated recognition, *Ry_sto_* and control tobacco leaves were site-inoculated with two other PPV isolates (PV-0001 and PV-2233 DSMZ). Western blot analysis did not detect PPV proteins in samples from the neighbouring leaf zones of *Ry_sto_* plants at 7 and 14 dpi whereas in non-transgenic control samples viral proteins were detected (Fig. 8b). To further validate Ry_sto_-mediated PPV resistance, we delivered a fulllength PPV cDNA clone tagged with GFP into transgenic *Ry_sto_* and control tobacco plants by agrofiltration. No GFP fluorescence was detected in the infiltrated leaves of *Ry_sto_* plants, whereas a strong signal was observed in control leaves (Fig. 8c).

**Fig. 8.**
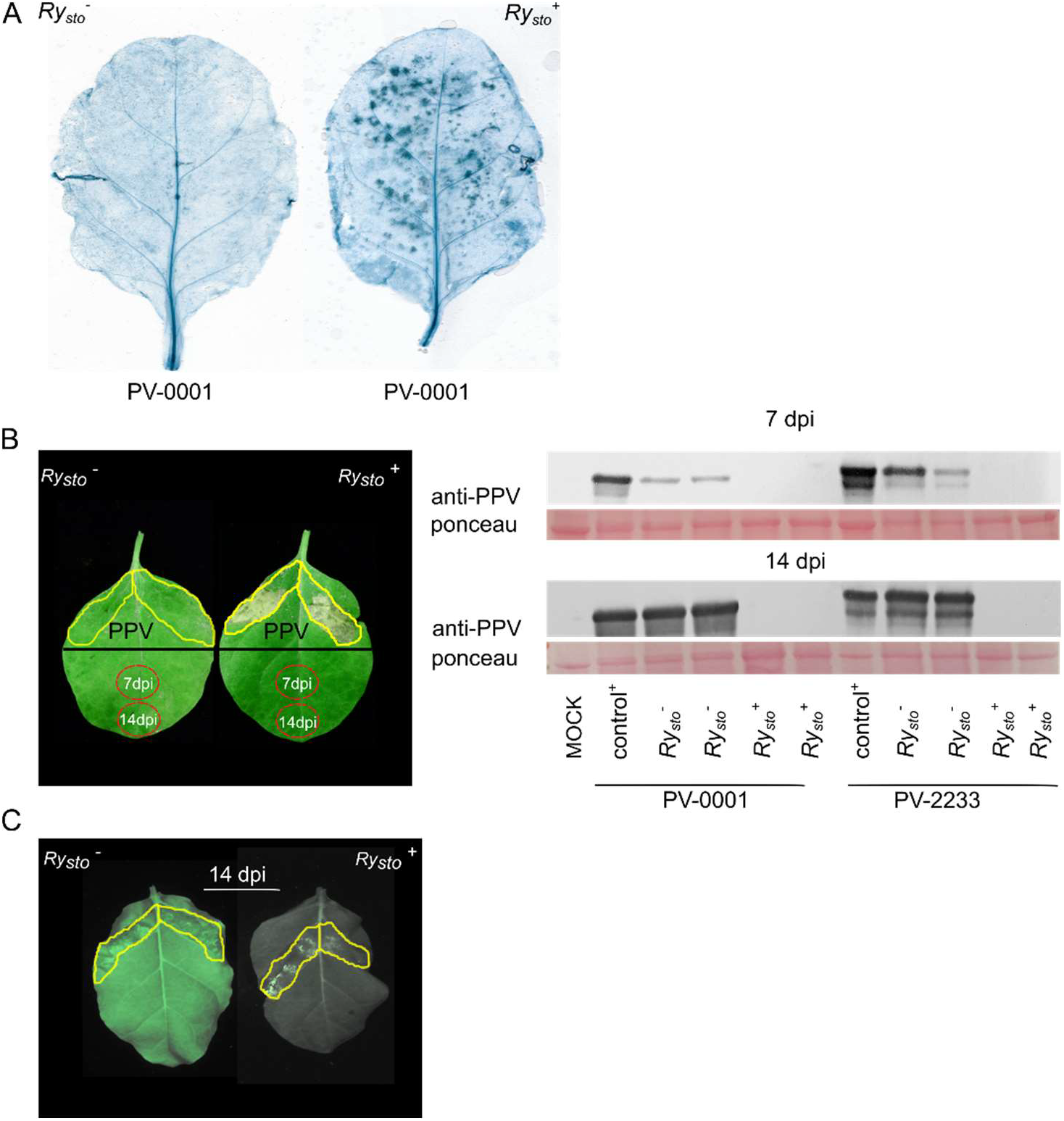
PPV triggers resistance response in *Ry_sto_*-expressing tobacco plants. (a) PPV-induced cell death development visualised by trypan blue staining in Ry_sto_-expressing or control, tobacco plants locally inoculated with a severe PPV isolate (PV-001, DSMZ). At 7 dpi, *Ry_sto_* plants exhibited a local cell death, whereas controls remained symptomless. (b) Ry_sto_ confers resistance to short-distance PPV movement. *Ry_sto_* transgenic and control plants were site-infected with different PPV isolates (PV-0001 or PV-2233, DSMZ). At 7 and 14 dpi, samples from consecutive leaf zones were harvested and PPV proteins were detected using anti-PPV antibodies. Staining of RuBisCo with Ponceau S was used as a loading control. (c) Leaves of or Ry_sto_-expressing and control plants were agroinfiltrated with PPV tagged with GFP. The viral spread was monitored by GFP-fluorescence. Photographs were obtained at 14 dpi under blue light (488 nm).

Unlike PPV, TuMV is able to infect *N. tabacum* systemically (Modarresi et al., 2019). Thus, after agroinfiltration of transgenic Ry_sto_ and control plants with a TuMV-GFP clone, GPF fluorescence and viral protein accumulation were monitored in upper, non-infiltrated leaves. At 21 dpi, no signal was detected in the Ry_sto_ plants, while control plants showed intense fluorescence (Fig. 9a). Similarly, western blot did not detect the viral proteins presence in transgenic *Ry_sto_* plants (Fig. 9b).

**Fig. 9.**
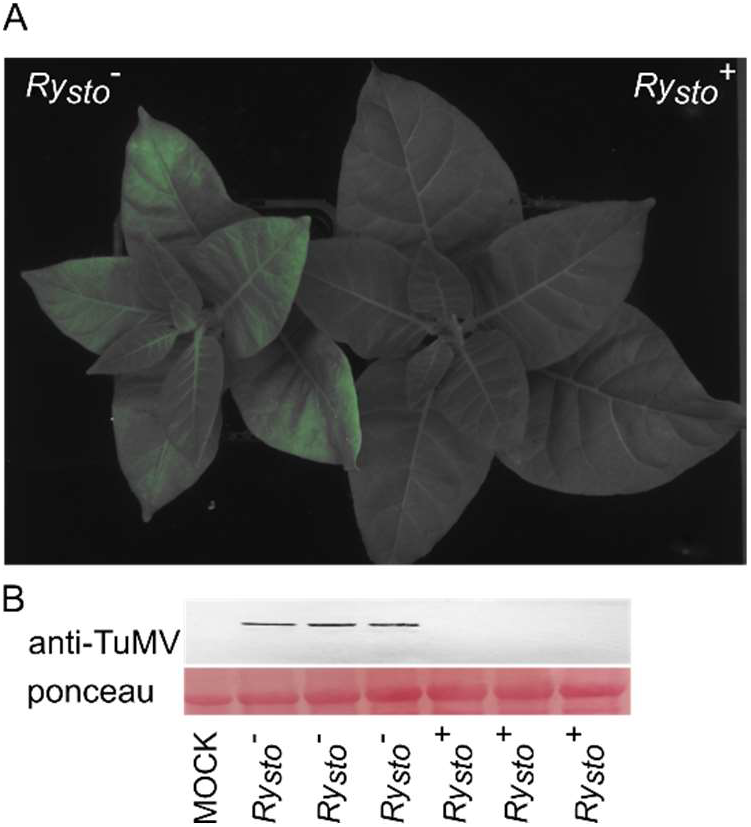
Ry_sto_ confers systemic resistance to turnip mosaic virus (TuMV). (a) Ry_sto_-expressing and control tobacco plants were infected by agroinfiltrated with TuMV - GFP. After 21 dpi control plants showed intensive GFP-fluorescence, while no signal was detected in Ry_sto_-expressing plants. The photograph was taken under blue light (488 nm). (b) SDS–PAGE analysis of samples from leaves of different tested plants was performed with anti-TuMV antibodies. Staining of RuBisCo with Ponceau S was used as a loading control.

Taken together, our results demonstrate that *Ry_sta_* immune receptor can trigger resistance against multiple potyviruses. We propose that Ry_sto_-CP interactions may represent one of the possible universal models of perception in which a single R receptor can recognise diverse Avr factors using a well-conserved element as a molecular target.

## Discussion

How NLRs sense pathogens and activate immunity is a fundamental question in plant immunity and disease resistance (Hu et al., 2020; Jones et al., 2016). The Ry_sto_ TIR-NLR receptor, which perceives conserved structural elements of the CPs of multiple potyviruses, may provide a useful model to understand how NLRs convert recognition to defence activation, and cope with the rapid evolution of viral pathogens.

Potyviruses are the second largest group of plant viruses, causing severe losses in many crops (Gadhave et al., 2020). Interestingly, they share a CP folding pattern that is widely distributed in eukaryotic viruses, including influenza viruses (Agirrezabala et al., 2015). As the CP features in virion assembly, RNA silencing and viral translation, it constitutes a potential universal target for R proteins (Zamora et al., 2015).

We identified the CP of PVY as the recognized component that triggers an Ry_sto_-mediated defence response in potato and other *Solanaceae* (Grech-Baran et al., 2020). Recent crystallographic studies of PVY CP revealed a disordered nature of N- and C-terminal ends and a conserved folding of its core (Kezar et al., 2019). Additionally, the N-terminal DAG motif was found to be indispensable for Potyviridae transmissibility (Revers et al., 2015). Our results provide evidence that only the 149 amino acid CP core region is required for Ry_st*o*_ perception, which is consistent with the preservation of its structure among the majority of potyviruses.

Many viral proteins execute multiple roles in the replication process, as well as in the suppression of plant defences (Leisner and Schoelz, 2018). In many cases, a combination of structural features creates defined motifs exerting particular functions (i.e., RNA binding), conserved in proteins from virtually all organisms (Navarro et al., 2020; Zamora et al., 2015). Such structure-function relationships are yet to be elucidated for many R-Avr interactions. It has been previously shown that a minimal fragment of 90 aa of PVX CP, encompassing four alpha helices, is sufficient to trigger Rx-mediated resistance (Nemykh et al., 2008). This suggests that the Rx sensing machinery targets conserved structural elements rather than particular sequence motifs (Baures et al., 2008). However, a single or double mutation in tobamoviral CP can disrupt *R*-gene-mediated resistance (Antignus et al., 2008), suggesting that certain residues and sequence motifs are crucial for triggering the response.

The S_125_, R_157_, and D_201_ residues that bind to ssRNA of potyviruses are crucial for proper tertiary folding of PVY CP (Zamora et al., 2017). Here, we showed that a single or double substitution of these amino acids impaired the cell death phenotype in *Ry_sto_* plants, whereas the triple mutant completely abolished it. The macroscopic HR phenotype was found to correlate with changes in the expression of defence-related markers, ion conductivity, and ROS production. Moreover, the mutations in the conserved R and D residues of the CP were shown to impair the *in vitro* assembly of the potyvirus johnsongrass mosaic virus (Jagadish et., 1993) and to block the assembly and cell-to-cell movement of the potexvirus pepino mosaic virus (Agirrezabala et al., 2015).

R_157_ and D_201_ residues form salt bridge bonds which contribute to the proper protein structure and specificity of interaction with other biomolecules. In general, disrupting salt bridges is energetically less favourable than reversing them, and may result in a decrease in protein stability (Bosshard et al., 2004). Remarkably, the potentially less stable CP derivative (S_125_A_R_157_D) was still able to trigger cell death, unlike the presumably more stable triple mutant variant (S_125_A_R_157_D_D_201_R) with a reversed salt bridge (Suppl. Fig 6). As both mutant proteins were expressed at similar levels, we propose that the phenotype of the triple mutant might be caused by compromised RNA binding, which in turn affects CP conformation, rather than the stability of the molecule.

Activated plant intracellular NLR receptors form oligomers that are required for immunity signalling that broadly resemble mammalian NLR inflammasome scaffolds (Martin et al., 2020; Ma et al., 2020). Two sub-families of helper NLRs, HeLo domain family (NRG1s and ADR1s) and EDS1 family of plant-specific lipase-like proteins (EDS1, PAD4 and SAG101) mediate defence signalling downstream of sensor TIR-NLRs (Lapin et al., 2019). Recently, it was reported that two distinct modules (NRG1/EDS1/SAG101 and ADR1/EDS1/PAD4) are required for TNLs-mediated immunity (Sun et al., 2020). Here, we provide evidence that Ry_sto_ mediates immune responses in EDS1-SAG101-NRG1 but not PAD4 dependent manner. A similar mechanism was described for another TNL, Roq1 (Sun et al., 2020).

Another major role of EDS1-PAD4 is to promote SA biosynthesis and activate SA-dependent defence responses (Gantner et al., 2019). We showed that Ry_sto_-mediated responses to transient CP expression are independent of SA accumulation, which is consistent with PAD4-independent function. Although transient CP expression in *Ry_sto_* plants was followed by some increase in SA levels, the cell death phenotype observed after the co-expression of Ry_sto_ with CP in *NahG* plants, suggests that the increase in SA is not required for activation of Ry_sto_-triggered response. Recently, it was reported that the NLR-dependent cell death response is strongly enhanced by the activation of surface receptors (Ngou et al., 2021). As viral coat proteins act as PAMPs (Teixeira et al., 2019), and also our Agrobacterium transient expression floods leaf apoplastic spaces with bacterial PAMPs, we cannot exclude that the overexpression of PVY CP boosts PTI and, therefore, the local increase in SA levels was independent of Ry_sto_ activation. Alternatively, the dying cells might release signals conditioning adjacent cells to become CP-responsive and activate SA biosynthesis and immunity throughout the entire plant (Radojičić et al., 2018).

Achieving complete and durable resistance is the ultimate goal of resistance breeding. We report here that the PVY CP core region architecture is highly conserved throughout the genus *Potyvirus*. Moreover, we show that the conserved core structure may serve as a rapid molecular marker for predicting the viral targets of Ry_sto_. Therefore, it is not surprising that a gene targeting such a conserved motif is both durable and broad-spectrum (Grech-Baran et al., 2020). Ry_sto_ recognises CPs of at least ten economically important potyviruses, including PRSV, PVM, SMV, PepSMV, BCMNV, MDMV, SCMV, TuMV, SPFMV, and PPV, which makes it a desirable candidate to engineer potyviral disease resistance into a range of crops worldwide.

To date, only a few sources of resistance to PPV have been identified, and no R genes were cloned so far (Ilardi et al., 2015; Zuriaga et al., 2018). We show here that Ry_sto_-expressing tobacco plants display resistance to both severe and mild PPV isolates, suggesting that *Ry_sto_* can be a valid alternative to RNAi-mediated resistance (Hily et al., 2007), provided that consumer resistance and regulatory constraints for GM crops were reduced. Similarly, several resistance genes (including *Tu*, *TuRB01*, *TuRB02*, *TuRB04*) were described for TuMV, but none were cloned so far, and several resistance-breaking strains were described (Palukaitis and Kim, 2021). Like *TuRB04* (Jenner et al., 2002), *Ry_sto_* confers extreme resistance to TuMV. Our *in silico* analysis, however, suggests that Ry_sto_ might resist TuRB04-breaking strains, thus providing a valuable additional candidate for resistance gene stacking to obtain truly broad-spectrum resistance.

Collectively, our study not only provides new insights into the mechanism of the *R*-gene surveillance system but also offers solutions for crop resistance against potyviruses, which are emerging as one of the most serious challenges of current agriculture, especially as the use of insecticides is becoming more restricted.

## Author contributions

M.G-B., and J.H performed most of the experimental work, data analyses and writing; K.W, J.P. performed bioinformatical analyses, T.M, A. GU performed the research; M.L performed confocal scanning, J.J. and K.W edited the article.

## Materials and Methods

### >Plant material

*Nicotiana tabacum* cv. Xanthi-nc, *Ry_sto_* transgenic *N. tabacum* and *Nicotiana benthamiana* plants were grown for 6 weeks in soil under controlled environmental conditions (22°C; 16 h light and 8 h dark) as described previously (Hoser et al., 2013). Transgenic tobacco plants were regenerated following *Agrobacterium tumefaciens* leaf disc transformation as described by Grech-Baran et al. (2020).

### Molecular Modelling

Each of the 10 selected CP proteins of *Potyviridae* were modelled by homology using the algorithm implemented in YASARA structure package (Krieger and Vriend, 2014). In all cases the same three template structures from Watermelon mosaic virus (5ODV), Turnip mosaic virus and (6T34) and Viruslike Particles based on Potato Virus Y (6HXZ) were selected. For each template, up to five alternative sequence alignments were allowed, and up to 50 different conformations were tested for each modified loop. Each of the obtained models was assessed for structural quality (dihedral distribution, backbone, and side-chain packing), and the ones with the highest scores were then used to create a hybrid model that was built using the best fragments (e.g. loops) identified in the particular models. The above procedure was then repeated for the folded parts of the proteins identified in the initial models. The structures of protein complexes with RNA were modelled using as the template two uridine pentanucleotides bound to Watermelon mosaic virus protein (5ODV, molecules a b and A, respectively). In the latter case the side-chain conformation was tuned by the ten succeeding rounds of FoldX 5.0 (https://doi.org/10.1093/bioinformatics/btz184) “repairPDB” procedure. Finally, the effect of all single residue replacement of RNA on the stability of the Protein-RNA complex was assessed using RNAscan procedure implemented in the FoldX 5.0.

The phylogenetic tree was generated from protein sequences of the known potyviruses obtained from NCBI. Sequences were aligned using ClustalW 1.7416 and the alignments were imported to the MEGA717 to build a maximum-likelihood phylogenetic tree with Jones-Taylor-Thornton (JTT) substitution model and 100 bootstraps.

### Plasmids generation

The sequences for all the primers used in this study are shown in Supplementary Table 1.

The PVY CP encoding cDNA was amplified by reverse transcription-PCR from total RNA extracted from PVY-infected tobacco leaf tissue as described earlier (Grech-Baran et al., 2020) and inserted into the pENTR/D-TOPO vector (Thermo Fisher Scientific, Waltham, US). The sequence of: Papaya ringspot virus (X67673 - 35K), Potato virus M (M96425.1), Soybean mosaic virus (GT015011), Pepper severe mosaic virus (NC_008393), Bean common mosaic necrosis virus (AB734777.1), Maize dwarf mosaic virus (AF395135.1), Sunflower chlorotic mottle virus (GU181199.1), Pea-seed born mosaic virus (HQ185577.1), Sweet potato feathery mottle virus (AF015540.1), Turnip mosaic virus (NC_002509.2), Cassava brown Streak virus (NC_014791.1) and Plum pox virus (AB576045.1) CPs were synthetized *in vitro* (Twist Bioscience, San Francisco, USA) and inserted into the pENTR/D-TOPO vector. The resulting entry clones were LR recombined with the Gateway pGWB 441, 445 or 411 destination vectors (Nakagawa et al., 2007). Side-directed mutagenesis was used for PVY CP single (CP S_125_A, CP R_157_D and CP D_201_R) as well as double (CP S_125_A/R_157_D, CP S_125_A/D_201_R) or triple (CP S_125_A/R_157_D/D_201_R) mutants generation. The presence of the mutation was confirmed by sequence analysis.

The *Ry_sto_* cDNA sequence was amplified from *Ry_sto_* tobacco plants and inserted into pDONR201 entry vector (Thermo Fisher Scientific) followed by an LR Gateway reaction into the pGWB441 destination vector (Nakagawa et al., 2007).

### Semi-quantitative RT-PCR

Total RNA was treated with DNase I (Thermo Fisher Scientific, Waltham, MA) and subjected to reverse transcription using a mix of random hexamers and oligo-dT primers and a Revert Aid first Strand cDNA Synthesis Kit (Thermo Fisher Scientific). Semiquantitative RT-PCR was performed using DreamTaq (Thermo Fisher Scientific) with 25 to 30 amplification cycles followed by electrophoresis on 2% agarose gel stained with ethidium bromide. Primer pairs used in the PCR reaction are listed in Suppl. Table 1.

### Gene expression analysis-RT-qPCR

Gene expression analysis via RT-qPCR was performed using a LightCycler480 instrument and LightCycler480 SYBR Green Master Kit reagents (Roche, Indiana, IN). Relative gene expression levels were determined using a standard curve method, and the value for each target gene was normalized against the mean expression values of two reference genes, *EF1* and *L23*, as described previously (Liu et al., 2012). Each sample was tested with four technical replicates and two dilutions. Primers for qPCR are provided in Suppl. Table 1.

### Ion Conductivity

At the indicated time points, eight leaf disks (1 cm diameter) were cut from infiltrated zones and floated abaxial side up on 5 ml milliQ water for 10 min at 18°C with gyratory agitation (50 rpm). The conductivity of the water was measured with a WTW InoLab Multi 9310 IDSCDM83 benchtop meter and expressed in μScm^-1^.

### Cell death scoring

*Agrobacterium tumefaciens* strain GV3101 carrying the appropriate constructs were suspended in infiltration buffer (10 mM MgCl2, 10 mM MES, pH 5.6, 150 mM acetosyringone) and mixed prior to infiltration at final OD600. Bacteria were infiltrated into leaves of ~4 weeks old *N. benthamiana* plants using a 1ml needleless syringe. At 5 days post infiltration (dpi), detached leaves were imaged under UV light (488nm) on the abaxial side, and scored using ImageJ software according to the scale presented in Suppl. Fig 9.

### O_2_^-^ detection

Superoxide anion concentrations were assayed spectrophotometrically as described by Krzymowska et al. (2007)

### Membrane integrity test

Membrane integrity was performed as described (Preethi et al., 2017).

### Western blotting

Agroinfiltrated leaves were collected, frozen and ground in liquid nitrogen. Total proteins were extracted as previously described (Grech-Baran et al., 2020). Homogenates were centrifuged at 21,000 g for 10 min, and the supernatants were collected. All tested samples were separated on 12.5% SDS-PAGE and subjected to immunoblot analysis using primary monoclonal Anti-Flag or, Anti-GFP antibodies (Abcam, Cambridge, UK) and alkaline phosphatase-conjugated anti-mouse secondary antibodies (Sigma Aldrich, St-Louis, USA). To detect PPV or TuMV, alkaline phosphatase-conjugated anti-PPV or anti-TuMV antibodies (Bioreba, Reinach, Switzerland) were used. Immunoblots were developed using a NBT/BCIP colourimetric detection kit (BioShop Inc., Canada).

### SA measurement

Free SA was extracted and quantified essentially as described by Malamy et al., (1990) and. HPLC assayed as described (Krzymowska et al., 2007).

### Transient expression assay

Agrobacterium GV3101 strains carrying plasmids expressing PVY CP, its variants or coat proteins of other *Potyviridae* were used to infiltrate *Ry_sto_* transgenic or control *N. tabacum*, as described previously (Grech-Baran et al., 2020). HR-related phenotypes were assayed 2–3 days after infiltration. Each experiment was performed at least twice and included at least three independent biological replicates.

### PPV and TuMV resistance assays

PPV isolates: 0001 (DSMZ No.:PPV-0001) or 2233 (DSMZ No.:PPV-0233) were inoculated, as recommended by German Collection of Microorganisms and Cell Cultures GmbH. Research (DSMZ). PPV-GFP and TuMV-GFP infections were perform as described (Fernández-Fernández et al. 2001; Garcia-Ruiz et al., 2010), respectively.

### Confocal laser scanning microscopy

Subcellular localization of the fusion proteins was evaluated using a Nikon C1 confocal system built on TE2000E and equipped with a 60 Plan-Apochromat oil immersion objective (Nikon Instruments B.V. Europe, Amsterdam, The Netherlands). GFP was excited with the 488 nm line from an argon ion laser, while RFP was excited by a 543 nm helium-neon laser and detected using the 605/75 nm barrier filter. Confocal images were analysed using free viewer EZ-C1 and ImageJ software.

### Statistical analysis

Statistical analyses were conducted using R 3.2.2 within R Studio 0.99.483. Technical replicates consisted of replicate readings from the same plant in the same experiment, whereas biological replicates consisted of measurements obtained from independent plants. Data were analysed using a following pipeline: data were assessed for their suitability for parametric analysis by testing for the normal distribution of the residuals using a Shapiro–Wilk test. If the data were suitable for conducting parametric tests, repeated measures ANOVA followed by Tukey’s HSD (*p* < 0.001**) test were performed.

## Acknowledgments

Research was supported by OPUS 14 (2017/27/B/NZ9/01803) grant from the National Science Centre, Poland to JH. MGB was partially supported by SONATA 15 (2019/35/D/NZ9/03565) grant from the National Science Centre, Poland. We would like to thank our co-workers, in particular dr Magdalena Krzymowska and dr Agnieszka I. Witek, for critical comments on the manuscript. Prof Jane Parker and Dr Johannes Stuttmann for sharing seeds of single *pad-4* and double *pad-4*, *sag101b N. benthamiana* knockout plants. Prof Juan Antonio Garcia Alvarez and Dr Véronique Decroocq for sharing PPV-GFP infectious clone. Prof James Carrington for shearing TuMV-GFP infectious clone. We would like to acknowledge Agata Gorczyca for the technical support.

