## Supplementary material for "The Ry_sto_ immune receptor recognizes a broadly conserved feature of potyviral coat proteins": Suplementary Figures

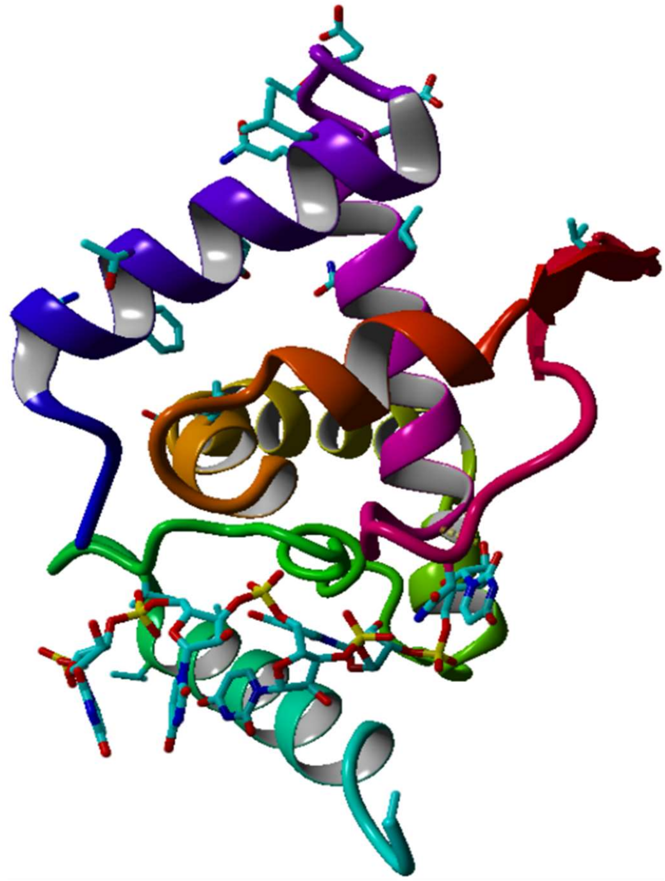

**Suppl. Fig. 1. Structural features of PVY.** Structure of the PVY CP unit modelled by homology. Disordered N- and C-terminal regions are omitted, whereas each of the seven α helices and one β hairpin is colour coded. The model was generated using the YASARA Structure package (Krieger and Vriend, 2014).

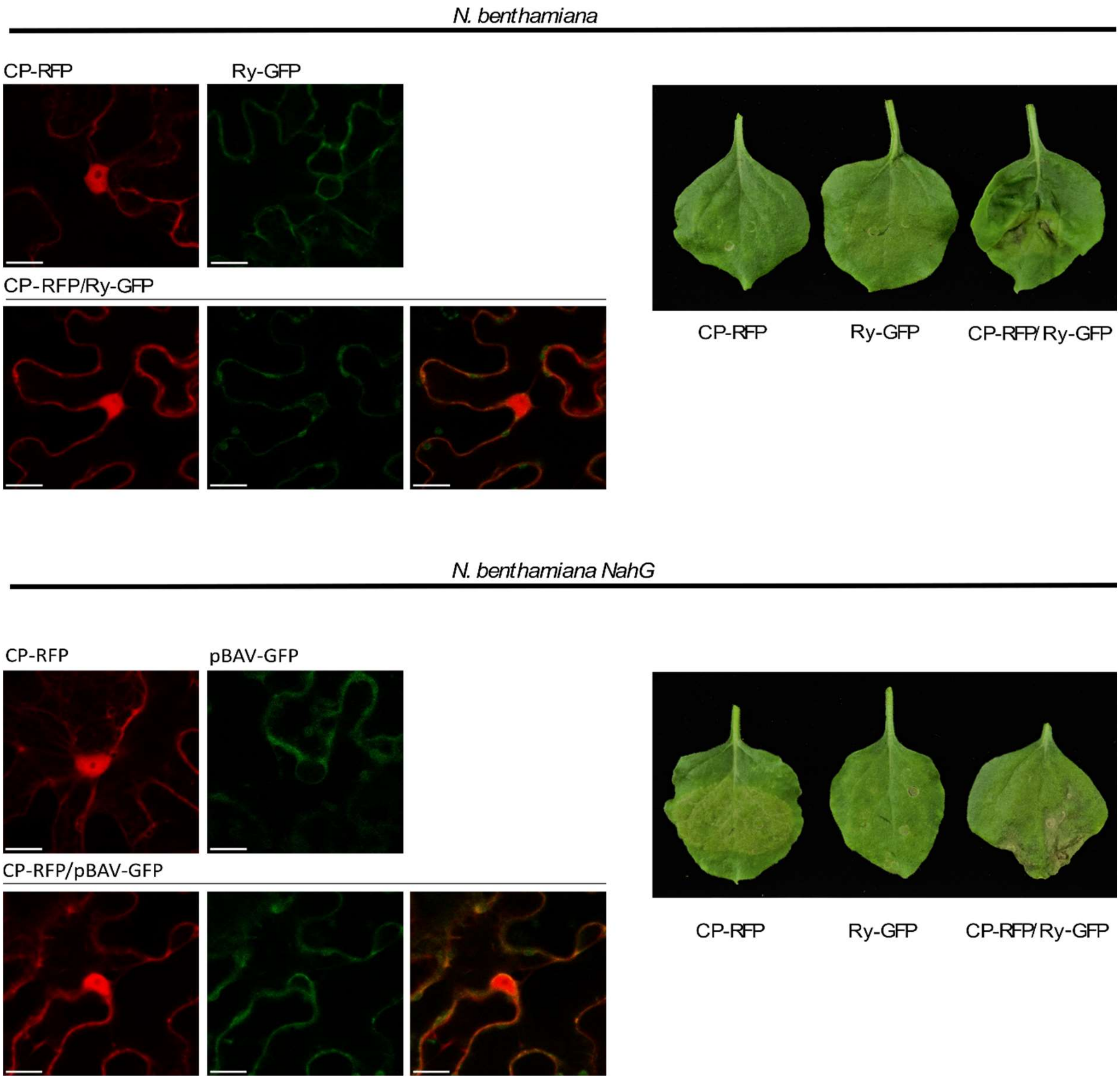


**Suppl. Fig. 2. Co-expression of Ry$_{sto}$ and CP results in SA-independent tissue collapse.** (right) Ry$_{sto}$-GFP, PVY CP-RFP or combination of both were expressed in leaves of control or *NahG N. benthamiana* plants using *Agrobacterium.* Twenty-four hours after infiltration (hpi), protein co-expression resulted in the development of cell death phenotype in both types of plants. The photograph was obtained at 72 hpi (left) Cellular localisation of the indicated proteins. Confocal images show representative leaf epidermal cells transiently expressing CP, Ry$_{sto}$, or both these proteins. Images were obtained 72 hpi. For each variant, approximately 50 transformed cells were examined. Bars = 10 µm

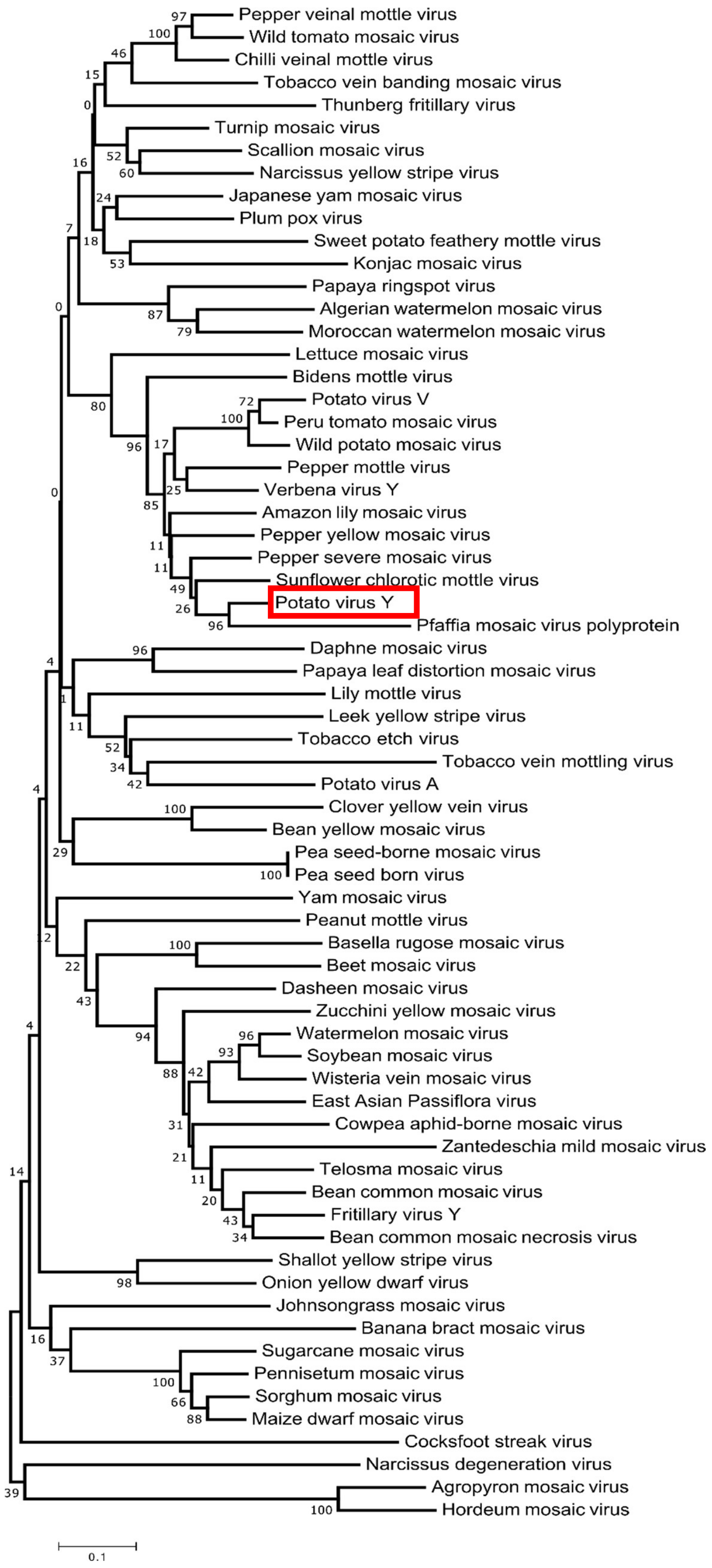


**Suppl. Fig. 3. Phylogenetic analysis of Potato virus Y CP protein and other *Potyviridae* CP proteins.** CP amino acid sequences were aligned using ClustalW and the alignments were imported to the MEGA717 to build a maximum-likelihood phylogenetic tree with Jones-Taylor-Thornton (JTT) substitution model and 100 bootstraps.

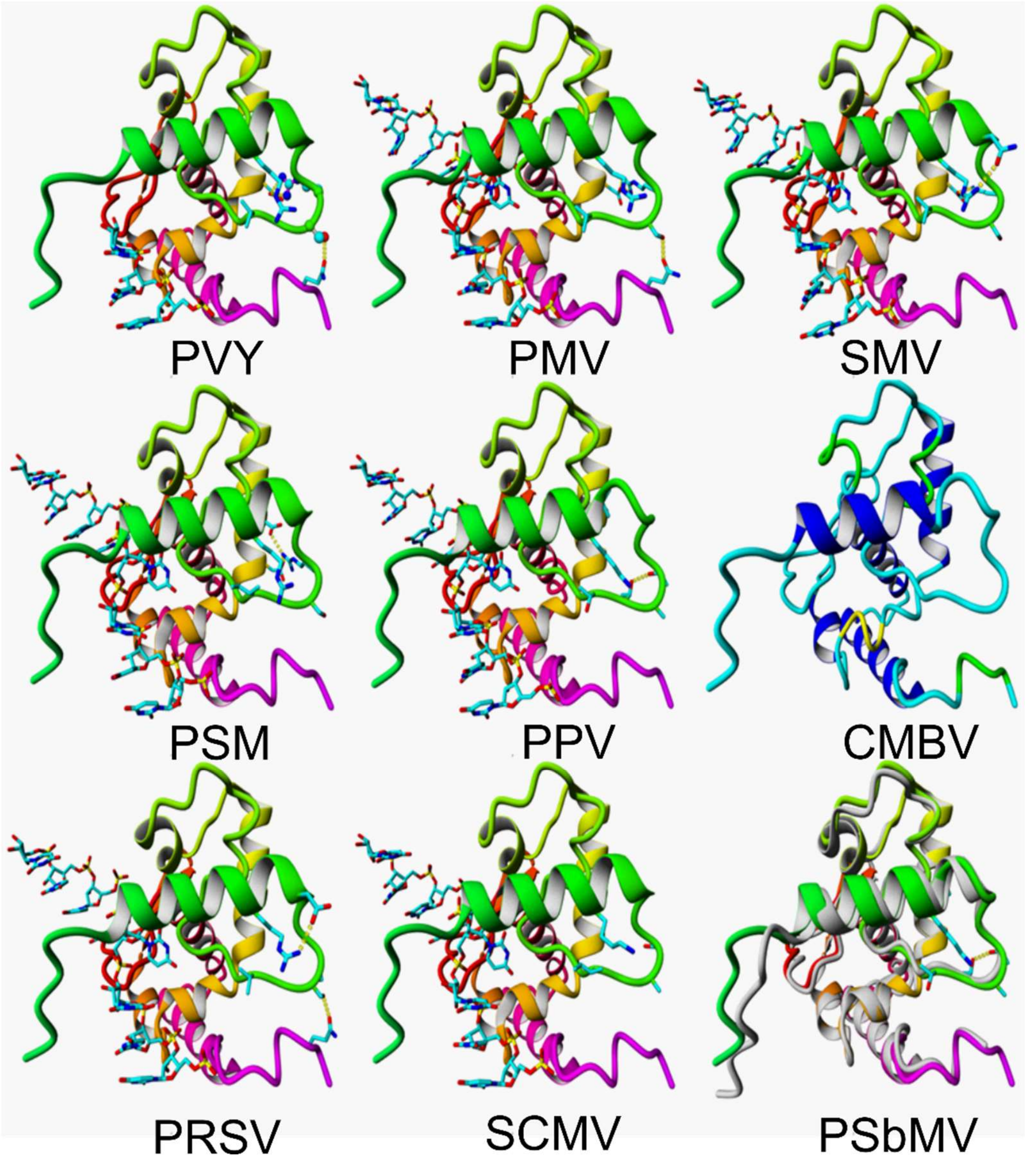


**Suppl. Fig.4. Structure of coat proteins of various potyviruses modelled by homology.** Models were visualised using Yasara Structure package (Krieger and Vriend, 2014).

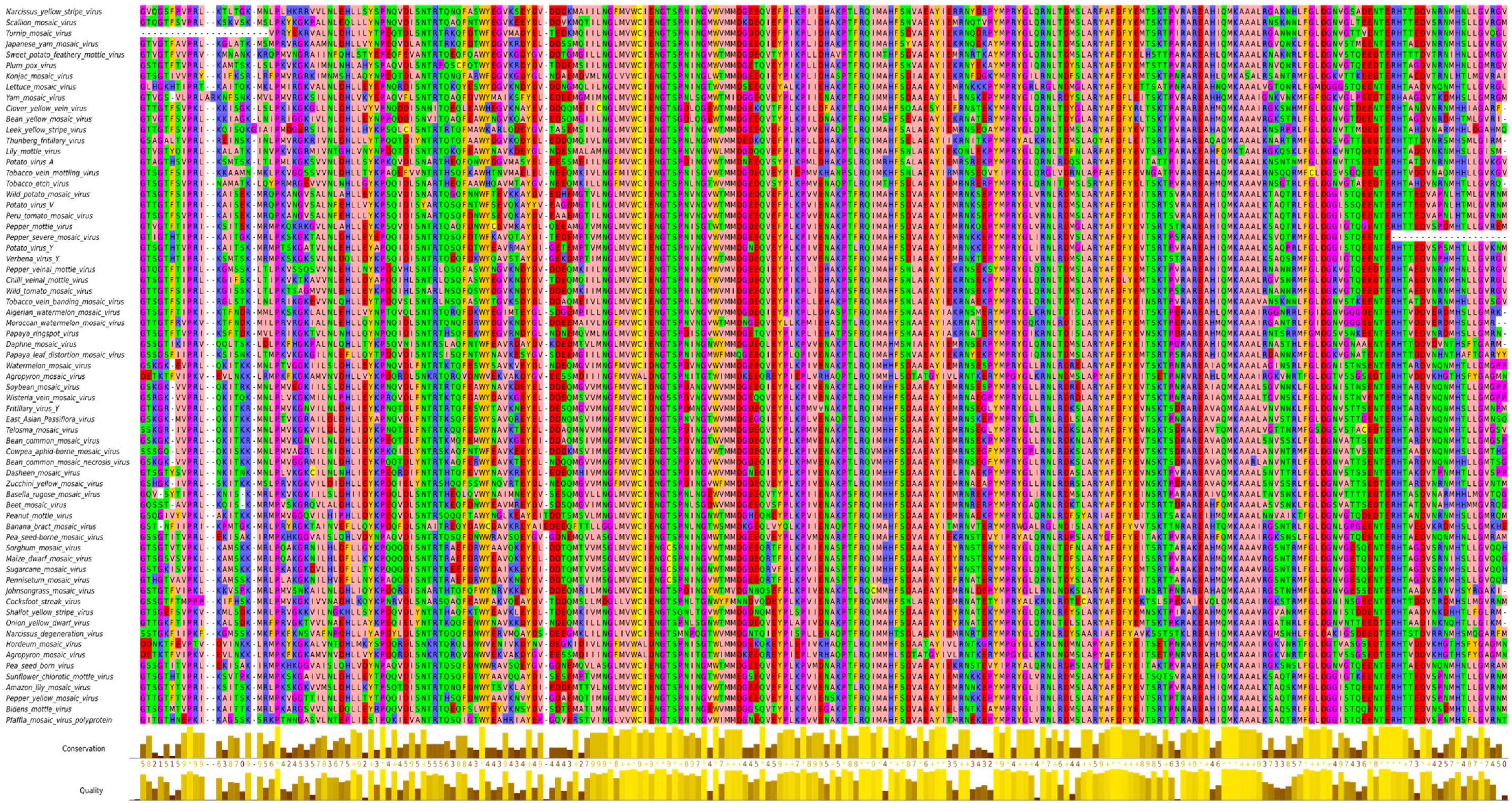

**Suppl. Fig. 5. Sequence alignment of conserved regions of potyviruses CP's core proteins.** The alignment was generated using MAFFT-L-INS-I (Katoh & Toh, 2008) and visualised in Jalview 2,10,4b1 (Waterhouse *et al.*, 2009)

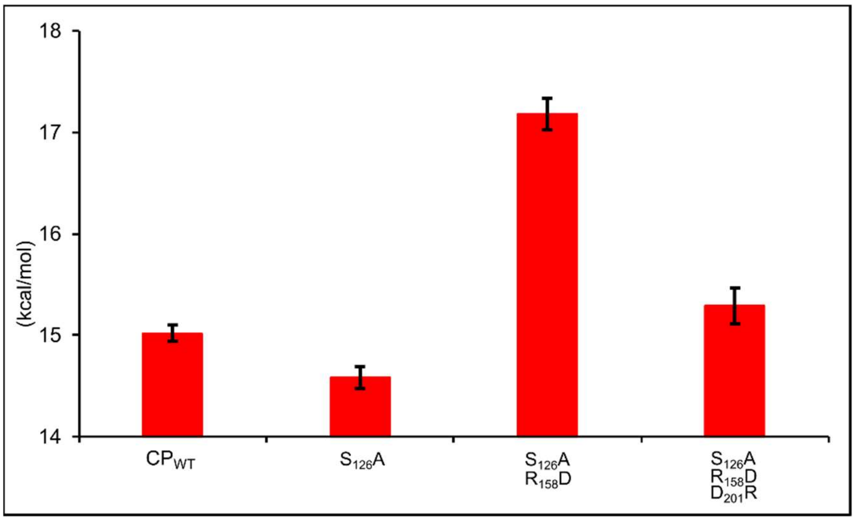


**Suppl. Fig. 6 Stability of CP variants (kcal/mol).** The effect of the single amino acids residue replacement on the stability of the CP-RNA complex was assessed using RNAscan procedure implemented in the FoldX 5.0.

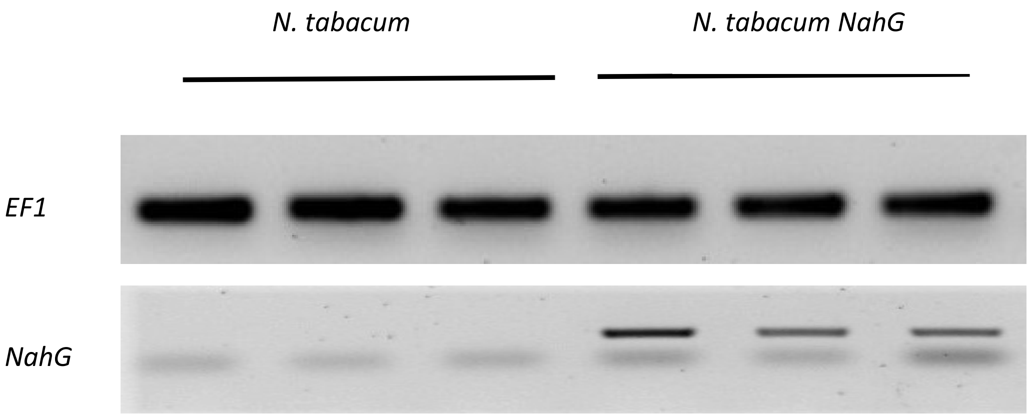


**Suppl. Fig. 7.** Semi-quantitative RT-PCR of *NahG* gene transcript. *N tabacum NahG* plants were obtained as described earlier (Friedrich et al., 1995). Elongation factor-1α (*EF1*α) was used as an internal control.

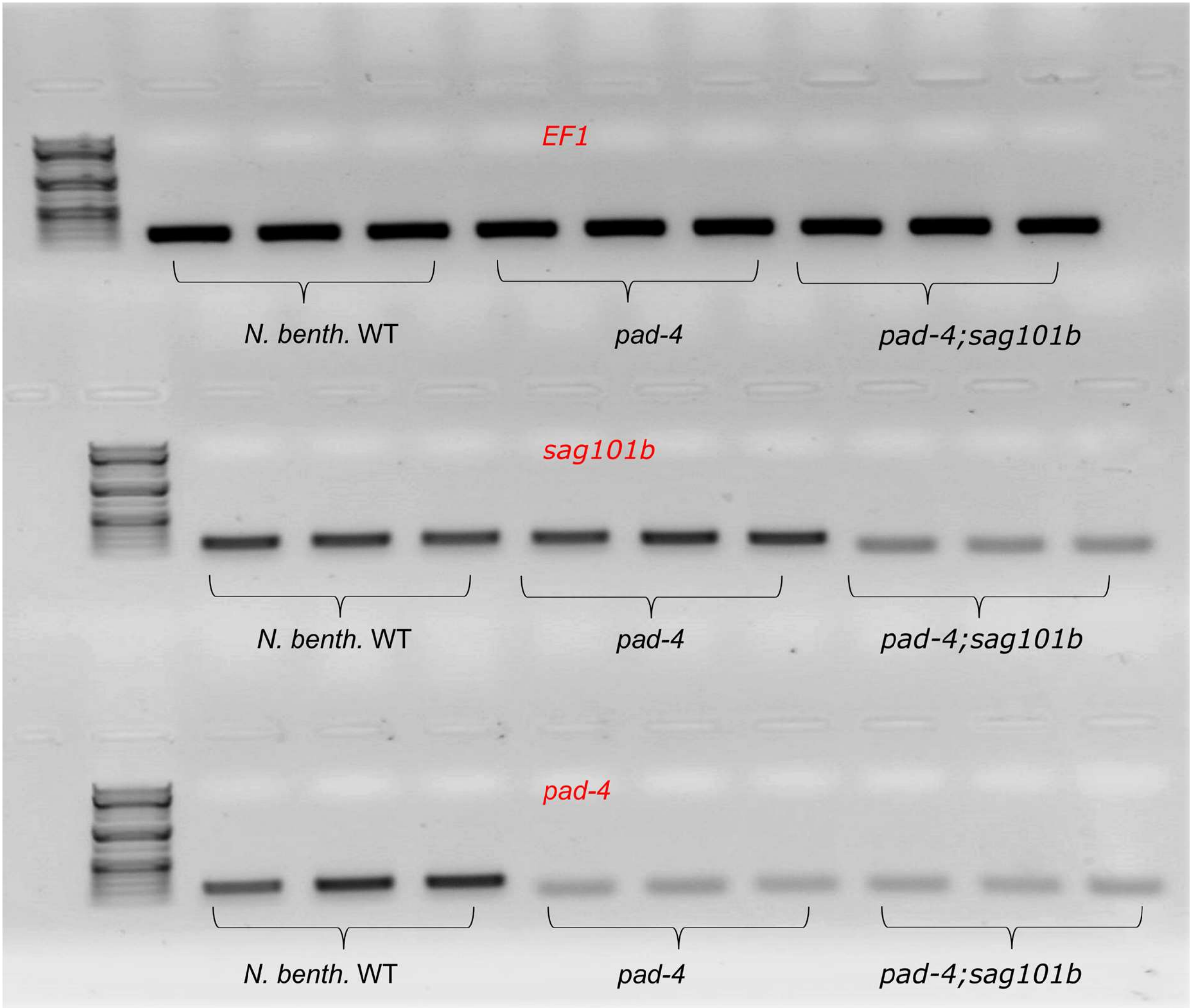


**Suppl. Fig. 8. Semi-quantitative RT-PCR of *Pad4 and SAG101b* gene transcripts.** Single *pad-4* and double *pad-4/sag101b N. benthamiana* knockout plants were made by Gantner et al., (2019). Elongation factor-1α (*EF1*α) was used as an internal control.

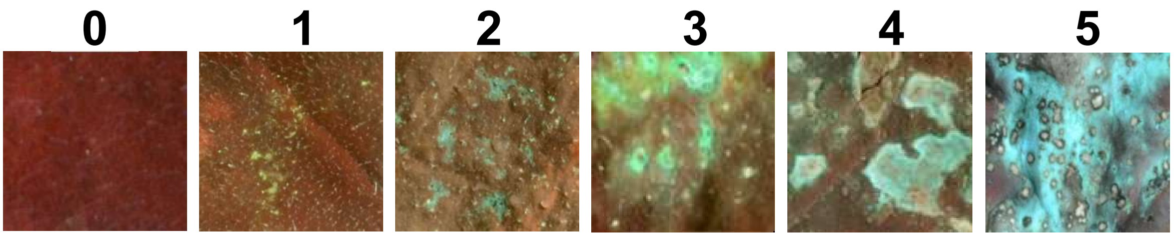


**Suppl. Fig. 9. Cell death rate scale.** Exemplary images used for scoring cell death index (HR Index) in *N. benthamiana* plants upon co-expression of $Ry_{sto}$ with CP, or CP mutants. Views from the adaxial side of the leaves for UV images were used.
