## Supplementary material for "The Ry_sto_ immune receptor recognizes a broadly conserved feature of potyviral coat proteins": Suplementary Tables

**Supplementary Table 1. Primers used to clone potyviral CPs.**

| ORF | Primer | Sequence |
|---|---|---|
| PVY CP | PVY_CP_F | **CACCATG**GGAAATGACACAATCGATGCAGGAGGAA |
| | PVY_CP_R | CATGTTCTTCACTCCAAGTAGAGTATGCATAC |
| PPV | PPV_CP_F | **CACCATG**GCTGATGAAAGAGAAGACGAGGA |
| | PPV_CP_R | CACTCCCCTCACACCGAGGAGGTTG |
| SPFMV | SPFMV_CP_F | **CACCATG**TCTAGTGAACGTACTGAATTCAAAGATG |
| | SPFMV_CP_R | TTGCACACCCCTCATTCCTAAGAGGTTA |
| CBMNV | CMV_F | **CACCATG**GAGAAAGATGCTGGAGCAGACAAA |
| | CMV_R | CTGTCCGGAAGTCATACCAAGTAA ATGATG C |
| MDMV | MDMV_F | **CACCATG**GCTGGTGAAAATGTTGACGCTGGAC |
| | MDMV_R | GTGTCCCTGCTGAACTCCCAGAAGG |
| PepSMV | P_SMV_F | **CACCATG**GGTGACAACATAGATGCAGGAAAAG |
| | P_SMV_R | CATGTTACGAACCCCAAGCAAAGTATGC |
| PbSbMV | PbSbMV_F | **CACCATG**GCTGGTGACGAAACCAAGGA |
| | PbSbMV_R | CATGGCTCTCATTCCGAGAAGATTGTG |
| PVM | PMV_F | **CACCATG**GCCGATACAACTGTTGATGCT |
| | PMV_R | TTATGTGTTTCTAACCCCAAGCAGAGTATG |
| PRSV | PRSV_F | **CACCATG**TCAAGGGGCACTGATGATTATC |
| | PRSV_R | TTATTAGTTGCGCATACCCAGGAGAG |
| SMV | SMV_F | **CACCATG**TCAGGCAAGGAGAAAGAAGG |
| | SMV_R | TTACTGCTGTGGGCCCATGCCCAGA |
| TuMV | TuMV_F | **CACCATG**GCAGGTGAAACGCTTGATGCAGG |
| | TuMV_R | CAACCCCTGAACGCCCAGTAAGTTAT |

In bold - extension for TOPO cloning

Normal font - gene specific sequence

**Supplementary Table 2. Primers used for site-directed mutagenesis of PVY CP**

| Gene | Primer | Sequence |
|---|---|---|
| CP_S126/A | CP_S_F | GGAACCGCGCCAAACATCAACGGA |
| | CP_S_R | TCCGTTGATGTTTGGCGCGGTTCC |
| CP_R157/D | | ACACTTGACCAAATCATGGCACATTC |
| | | AATGTGCCATGATTTGGTCAAGTGT |
| CP_D201/R | | GTAACTTCATAAAAGCGAAAAGCATA |
| | | TATGCTTTTCGCTTTTATGAAGTTAC |

**Supplementary Table 3. Primers used for $Ry_{sto}$ cDNA cloning**

| Gene | Primer | Sequence |
|---|---|---|
| Ry | Ry_F | **GGGGACAAGTTTGTACAAAAAAGCAGGCTACACC**ATGCTTCTTCTTCC |
| | Ry_R | **GGGGACCACTTTGTACAAGAAAGCTGGGTC**ACATTTCATATATAAGATAAAAACCTGG |

In bold – an extension for pDONR cloning

Normal font – a gene specific sequence

**Supplementary Table 4. Primers used for qPCR and RT PCR reactions**

| Gene | Primer | Sequence |
|---|---|---|
| *EF1* | EF1_F | **GACAAGCGTGTTATTGAGAGG** |
| | EF1_R | CACAGTGCAGTAGTACTTAGTG |
| *HSR203_J* | HSR_F | GGGAACATAGTCCACCAAGTC |
| | HSR_R | ACTCCGATTTGCTCCGATAAG |
| *HIN-1* | HIN_F | ATCCTCGGAGTCATTGCATTAG |
| | HIN_R | TGTTGTTTGTGGTGGACAAATC |
| *L23* | L23_F | AAGGATGCCGTGAAGATGT |
| | L23_R | GCATCGTAGTGGAGTCAAC |
| *NahG* | NahG_F | AGGGCGGCGCAGGTCGTCGTAGGCTTCA |
| | NahG_R | GGCGGCATCATCAACGTGGTGGCTTTCA |
| *PAD-4* | PAD-4_F | GCGCTGGAGGAGTCGTGGAAGGCTTGC |
| | PAD-4_R | TCGGGAAGCGGCAACAAATTCCGGCAAC |
| *SAG101b* | SAG101b_F | CAGGATTCCTTGTGAGTGTGGCTATCT |
| | SAG101b_R | CACCACCAACTTTGCCAATTCTTGGCCAC |
